# Rapid toxin sequestration modifies poison frog physiology

**DOI:** 10.1101/2020.05.27.119081

**Authors:** Lauren A. O’Connell, LS50: Integrated Science Laboratory Course, Jeremy D. O’Connell, Joao A. Paulo, Sunia A. Trauger, Steven P. Gygi, Andrew W. Murray

## Abstract

Poison frogs sequester chemical defenses from their diet of leaf litter arthropods for defense against predation. Little is known about the physiological adaptations that confer this unusual bioaccumulation ability. We conducted an alkaloid-feeding experiment with the Diablito poison frog (*Oophaga sylvatica*) to determine how quickly alkaloids are accumulated and how toxins modify frog physiology using quantitative proteomics. Diablito frogs rapidly accumulated the alkaloid decahydroquinoline within four days, and dietary alkaloid exposure altered protein abundance in the intestines, liver, and skin. Many proteins that increased in abundance with decahydroquinoline accumulation are plasma glycoproteins, including the complement system and the toxin-binding protein saxiphilin. Other protein classes that change in abundance with decahydroquinoline accumulation are membrane proteins involved in small molecule transport and metabolism. Overall, this work shows poison frogs can rapidly accumulate alkaloids, which alter carrier protein abundance, initiate an immune response, and alter small molecule transport and metabolism dynamics across tissues.

**Summary Statement:** Poison frogs rapidly accumulate toxins, which changes abundance of proteins involved in the immune system and small molecule binding and metabolism across tissues.

## Introduction

Many organisms can acquire chemical defenses from external sources. These acquired chemical defenses are sometimes diet-derived and many examples can be found in invertebrates (Opitz and Müller, 2009) and less commonly in vertebrates (Savitzky et al., 2012). Poison frogs sequester lipophilic alkaloids from arthropod prey (Saporito et al., 2009), which requires specialized physiology for bioaccumulation and toxin resistance (Santos et al., 2016). The bioaccumulation of alkaloids requires coordination between several tissue systems, from absorption in the gut, transport between tissues, and storage in skin granular glands. Although sequestering defensive chemicals involves a well-studied ecological relationship between arthropod prey and frog predators (Darst et al., 2005; Saporito et al., 2011), how poison frogs sequester alkaloids is unknown.

The chemical defenses of poison frogs are mainly lipophilic alkaloids and over 800 have been characterized in poisonous amphibians (Daly et al., 2005). There is exceptional species diversity in chemical defenses, particularly in South American dendrobatid poison frogs (Saporito et al., 2011). The ability to sequester alkaloids evolved several times in dendrobatids and coincides with a dietary specialization on ants and mites (Darst et al., 2005; Santos et al., 2003). The dietary hypothesis of poison frog toxicity arose when several studies noted that poison frogs reared in captivity lacked alkaloids (Daly et al., 1994a) and subsequently examined alkaloid uptake (Daly et al., 1994b; Hantak et al., 2013). Research over several decades has focused on alkaloid characterization (Saporito et al., 2011), but how poison frogs accumulate toxins remains largely unexplored.

Although alkaloids are lipophilic, these small molecules do not simply perfuse passively throughout the organism. Evidence for an uptake system in poison frogs comes from alkaloid quantification in various tissues, including the liver, muscles, and oocytes (Prates et al., 2011; Stynoski et al., 2014), although alkaloids are most abundant in the skin where they are stored in granular glands (Neuwirth et al., 1979). Moreover, the vertebrate intestinal lining is well-equipped to prevent passive absorption of toxic substances using a series of membrane transporters (Zhang and Benet, 2001). Absorbed lipophilic molecules are either transported through the lymph system by chylomicrons or through the blood circulation bound to carrier proteins produced by the liver. The molecular physiology of alkaloid bioaccumulation, metabolism, and transport between each of these poison frog tissue systems is unknown.

We conducted an experiment to determine how alkaloid sequestration alters frog physiology and tested the general hypothesis that alkaloid bioaccumulation alters the abundance of proteins involved in small molecule transport and metabolism. We used the Diablito frog (*Oophaga sylvatica*) to address this question, which complements our previous studies examining the chemical ecology of this species (Caty et al., 2019; McGugan et al., 2016; Roland et al., 2016). Frogs were fed the alkaloid decahydroquinoline (DHQ), a commercially available alkaloid they naturally sequester in the wild (Caty et al., 2019), and we examined how alkaloid sequestration influences frog physiology using quantitative proteomics. Our experiments show that poison frogs sequester alkaloids within four days of exposure, shifting protein abundance across tissues, including plasma glycoproteins and cytochrome P450s.

## Materials and Methods

### Animals

Adult Diablito frogs (*Oophaga sylvatica*, N=12, 5 females, all of unknown age) were purchased from Indoor Ecosystems (Whitehouse, OH, USA) and housed for three months prior to the experiment. Frogs were housed individually on a 12:12 hour photoperiod, immediately preceded and followed by a 10 min period of dim incandescent lights to simulate dawn and dusk. Frogs were misted with water three times daily and fed four times per week with fruit flies dusted with a vitamin supplement. The Institutional Animal Care and Use Committee of Harvard University approved all procedures (Protocols 15-02-233 and 15-07-247).

### Decahydroquinoline feeding experiment

Frogs were randomly assigned to one of four groups to total N=3 per group. Within-group variation was unknown during experimental design and an N=3 is standard in the quantitative proteomics field. The three experimental groups received fruit flies dusted with a 1% decahydroquinoline (DHQ; Sigma-Aldrich, St. Louis, MO, USA) in a vitamin mix (Dendrocare vitamin/mineral powder, Black Jungle, Turners Falls, MA, USA) for either 1, 4, or 18 days (Figure 1A). DHQ-like alkaloids are sequestered by *O. sylvatica* in the wild (Caty et al., 2019) and represent an ecologically relevant treatment. Control frogs (N=3) received flies dusted with vitamin mix without DHQ. For the 1-day group, frogs received toxin-dusted flies in the morning, and then were sacrificed the following day after receiving another dose of flies that morning (a total of 2 feedings in 36 hours). For the 4-day group, frogs were fed toxic flies on Monday, Tuesday, Thursday, and Friday mornings and were sacrificed Friday afternoon. For the 18-day group, frogs were fed toxic flies 4 days per week for 3 weeks for a total of 12 feedings. Roughly 15-25 flies (average = 18) flies were given to the frogs at each feeding session and the number of flies remaining the next morning were recorded. The average number of total alkaloid-dusted flies eaten by each group during the duration of the experiment were 14.3 for 1-day, 52.3 for 4-day, and 149.7 for 18-day. As DHQ was administered by dusting prey items, and since individual frogs ate a variable number of prey, we were unable to calculate the exact DHQ dose for each individual. On the afternoon of the last day, the frogs were anesthetized by applying 20% benzocaine gel to the ventral side and then euthanized by decapitation. Dorsal skin, liver, and intestines were dissected, flash frozen in liquid nitrogen and stored at −80°C for later processing.

**Figure 1.**
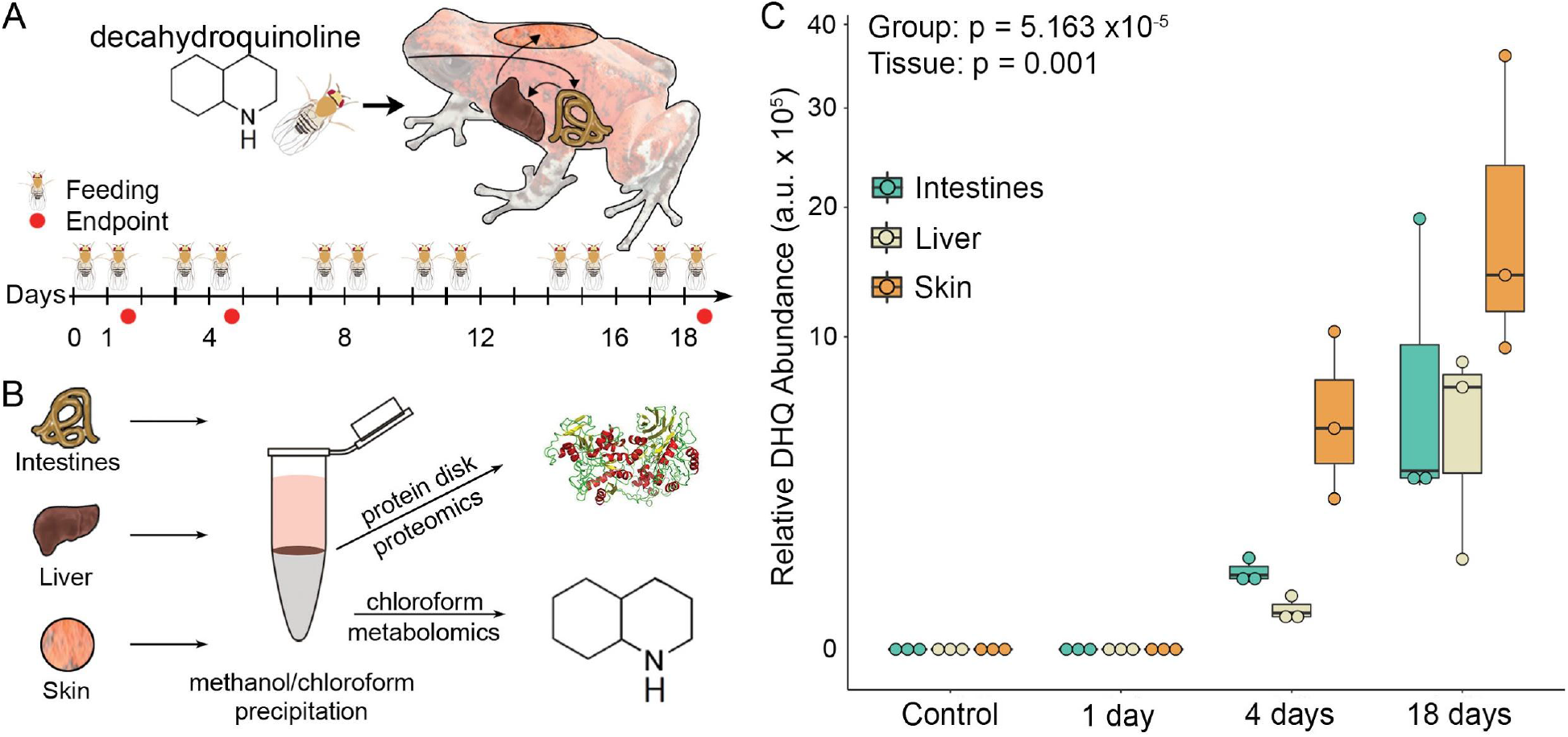
Rapid uptake of decahydroquinoline (DHQ). **(A)** Frogs were fed DHQ-dusted flies to determine the rate of toxin bioaccumulation (N=3 per group). The timeline shows the number of times that frogs were fed with toxic flies and the endpoints. This experiment was performed once in a teaching laboratory by undergraduate students. **(B)** Tissues were dissected and both protein and DHQ were isolated using methanol/chloroform precipitation. **(C)** DHQ abundance (y-axis, log scale) in the intestines (light green), liver (cream), and skin (light orange) increase over time, with the first detectable amounts within 4 days. Data are represented in boxplots that show rectangles as the lower and upper quartiles (with the median as the line) and whiskers that indicate the maximum and minimum values. A generalized linear mixed model showed a significant effect of group (p=5.163×10^−5^) and tissue (p=0.001).

To isolate proteins and alkaloids from the same tissue samples, flash frozen tissues were removed from −80°C and immediately crushed to a powder in liquid nitrogen using a mortar and pestle. Tissue powder was then resuspended in a lysis buffer from the Mammalian Cell Lysis kit containing a protease inhibitor cocktail (Sigma-Aldrich). Samples were incubated in lysis buffer at 4°C for 15 min rotating and then centrifuged for 20,000 x g for 10 min at 4°C to pellet cell debris. The supernatant was transferred to a chilled microcentrifuge tube and protein concentration was determined using 2 μl of lysate in a Bicinchoninic Acid (BCA) Assay according to manufacturer’s instructions (Thermo Scientific, Waltham, MA, USA). To standardize the amount of input material in downstream assays, lysate containing 300 μg of protein from each tissue type was transferred to a chilled microfuge tube for protein and alkaloid (DHQ) extraction. A chloroform-methanol extraction was used to separate the DHQ and proteins from the same tissue lysate sample. Briefly, 400 μl 100% methanol was added to each sample and then each sample was vortexed. Then 100 μl chloroform was added to each sample, which was immediately vortexed. To precipitate protein, 200 μl nuclease-free water was added to each sample. After vortexing, samples were centrifuged at 21,000 x g for 3 min at room temperature, resulting in a protein disk at the interface of the aqueous methanol top layer and the organic chloroform bottom layer. A previous pilot experiment showed roughly 90% of DHQ spiked into a protein lysate sample separated into the chloroform layer. Therefore, the methanol layer was removed and the chloroform layer was isolated into a glass vial and stored at −20°C for later quantification of DHQ. The remaining protein pellet was rinsed twice with 1 mL methanol and then stored at −80°C for further processing.

### Quantification of DHQ

Liquid chromatography / mass spectrometry (LC/MS) quantification of DHQ was performed on a Bruker maXis Impact Q-TOF system (Billerica, MA) with an Agilent 1290 LC (Palo Alto, CA). A reversed-phase LC gradient method was used using a Dikma C8, 3 μm particle size, 2.1 x 100 mm column (Dikma, Lake Forest, CA). Mobile phase A was composed of water with 0.1% formic acid and mobile phase B was composed of acetonitrile with 0.1% formic acid. The flow rate was 0.3 mL/min. The gradient began with 0% B for two min then increased linearly to 100% B at 8 min and was held until 10 min. The solvent composition was then returned to starting aqueous conditions to re-equilibrate the column for the next injection. The mass spectrometer was tuned for standard mass range analysis and data were continually acquired in the range m/z 50-3000; each run was recalibrated for the m/z scale using a post-run injection of a sodium formate solution. Electrospray positive mode ionization was used with a source drying gas of 10 L/min at 200°C, nebulizer at 30 psi, and capillary set at 4000 V with an endplate offset of 500 V. For the Agilent Q-TOF, the Ion Funnel electrospray positive mode source used drying gas of 14 L/min at 200°C with nebulizer at 35 psi, a sheath gas flow of 11 L/min at 350°C, capillary set at 3500 V, nozzle voltage of 1000 V, and Fragmentor set at 175 V. Collision energies were set at 15 and 30 eV and data were continually acquired in the range m/z 50-1700 using a reference lock mass. A DHQ standard was run along with the experimental samples to identify the correct peak and the structure was confirmed by its fragmentation pattern using tandem mass spectrometry (Figure S1).

### Quantitative proteomics

Protein disks were thawed and resuspended in 10 μl acetonitrile with gentle agitation to break up the disk, followed by the addition of 90 μl 8M urea in 100mM EPPS pH 8.5 with another round of gentle vortexing. Samples were spun down at 13,000g for 30s, and sonicated in a water bath for 20 min at room temperature to complete resuspension. Proteins disulfide bonds were reduced with 5 mM tris-(2-carboxyethyl)-phosphine (TCEP), at room temperature for 25 min, and alkylated with 10 mM iodoacetamide at room temperature for 30 min in the dark. Excess iodoacetamide was quenched with 15 mM dithiothreitol at room temperature for 15 min in the dark. Then a methanol-chloroform precipitation was performed again as described above. Protein disks were resuspended in 10ul of 8M urea, 200mM EPPS pH 8.5 with gentle vortexing followed by the addition of 90ul 200mM EPPS pH 8.5 to reach 100 μl and dilute to 0.8M urea. Proteins were then digested at room temperature with gentle agitation for 12 hrs with LysC protease at a 100:1 protein-to-protease ratio. Then, trypsin was added at a 100:1 protein-to-protease ratio and the reaction temperature raised to 37°C for 6 hrs. Digests were spun at 10,000 g for 10 min and the supernatant transferred to a new tube. Peptide concentrations were measured using the Quantitative Colorimetric Peptide assay kit (Thermo Scientific) according to the manufacturer’s instructions. Following quantification, 50 μg of peptides from each sample were labelled with 5 μl of Tandem Mass Tag 10-plex (TMT, Thermo Scientific) reagent (0.02mg/μl) in 200mM EPPS pH8.5 at 30% acetonitrile (v/v) for 1 hour before quenching the reaction with 10 μl of 5% hydroxylamine. A pool of each control sample was labelled collectively to create a bridge in an additional channel.

The TMT-labelled samples from each tissue from a given collection were then pooled in equimolar ratios, as verified by an MS2-only ratio check of an aliquot. The pooled sample was vacuum centrifuged to near dryness before resuspension in bicarbonate buffer, and separation by basic pH RP HPLC on an Agilent 1100 pump equipped with a degasser and a photodiode array (PDA) detector measuring at 220 and 280 nm wavelengths (Thermo Fisher Scientific - Waltham, MA). Peptides were subjected to a 50 min linear gradient from 8% to 80% acetonitrile in 10mM ammonium bicarbonate pH 8 at a flow rate of 0.6 mL/min over an Agilent 300Extend C18 column for 75 minutes (3.5 μm particles, 4.6 mm ID and 250 mm in length) to produce 96 equal volume fractions. The fractions were re-pooled down to 24 fractions, and vacuum centrifuged to reduce volume. The pooled fractions were acidified and then desalted via StageTip, vacuum centrifuged to near dryness, and reconstituted in 5% acetonitrile, 5% formic acid for injection into an Orbitrap Fusion Lumos. Twelve non-adjacent pooled fractions were analyzed via mass spectrometry.

Each fraction was separated along a 4-30% acetonitrile linear gradient over 3 hours, and ionized via electrospray for analysis by mass spectrometry using a TOP10 method, where each FTMS1 scan was used to select up to 10 MS2 precursors for CID-MS2 followed by measurement in the ion trap. Each MS2 was used to select precursors (SPS ions) for the MS3 scan which measured reporter ion abundance for the 10 samples simultaneously (McAlister et al., 2014). Instrument parameter settings included an FTMS1 resolution of 120,000, ITMS2 isolation window of 0.4 m/z,120 ms ITMS2 max ion time, ITMS2 AGC set to 2E4, ITMS2 CID energy of 35%, SPS ion count allowing up to 10 precursors, FTMS3 isolation window of 1.2 m/z, 150 ms FTMS3 max ion time, FTMS3 AGC set to 1.5E5, and FTMS3 resolution of 50,000.

Samples were searched with the Sequest algorithm (Ver. 28) against a custom *O. sylvatica* proteome database. There are no high quality genomes available for poison frogs and thus we are limited to transcriptome-based references. We confirmed this transcriptome-based method was the best approach by comparing the number of unique peptides that matched the *O. sylvatica* transcriptome-derived proteome compared to *Xenopus* and human reference proteomes (Figure S2). All analyses were performed with a database generated from a *O. sylvatica* transcriptome (Caty et al., 2019) using the PHROG workflow (proteomic reference with heterogeneous RNA omitting the genome; (Wühr et al., 2014)), which was concatenated with their reversed sequences as decoys for FDR determination. Common contaminant sequences were similarly included. For searches we restricted the precursor ion tolerance to 50 ppm, and set the product ion tolerance window to 0.9 m/z, allowed up to two missed cleavages, included static medication of lysine residues and peptide N-termini with TMT tags (+229.163 Da), static carbamidomethylation of cysteine residues (+57.021 Da), and variable oxidation of methionine residues (+15.995 Da). The search results were combined and filtered to a 1% FDR at the peptide and protein levels using linear discriminant analysis and the target-decoy strategy (Elias and Gygi, 2010; Huttlin et al., 2010). MS3 spectra were processed as signal-to-noise ratios for each reporter ion based on noise levels within a 25 Th window. Proteins quantitation was the result of summing reporter ion intensities across all matching PSMs. PSMs with isolation specificity below 0.7, MS3 spectra with more than eight TMT reporter ion channels missing, or MS3 spectra with a TMT reporter ion summed signal to noise ratio that is less than 200 were removed before analysis. Overall, several thousand proteins were detected in each tissue type (5328 in the intestines, 5837 in the liver, and 5987 in the skin).

### Data analysis

All statistics and figures were generated in R Studio (version 1.1.442) running R (version 3.5.2). For DHQ abundance, the integrated area under the peak from the ion chromatograph was used for sample quantification compared to the standard and values are presented in arbitrary units. We used the glmmTMB R package (Brooks et al., 2017) to run a generalized linear mixed model with a zero-inflated negative binomial distribution to test for significant differences in DHQ quantity between groups and tissue type as main effects. Frog identity was included as a random effect to account for the sampling of three tissues (intestines, liver, and skin) for each frog. We then followed the model with the Anova.glmmTMB function for reported statistical values. Boxplots were generated with the ggplot function in the ggplot2 package in R (version 3.3.0 (Wickham, 2009)).

For the proteomics data, protein contigs with fewer than two peptides were removed from the analysis, leaving 3715 in the intestines (69.7%), 3789 in the liver (64.9%), 3755 in the skin (62.7%). We combined the day 4 and day 18 animals with DHQ in our proteomics analysis, totalling N=6 in the “toxic” group and N=3 in the “non-toxic” or control group. Day 1 was not included as we did not detect any DHQ in those samples and tandem mass tags for labeling was, at the time, limited to 10 tags. We used the eBayes() function in the R limma package (Ritchie et al., 2015) (version 3.36.5) to determine significant differences between groups in each tissue (D’Angelo et al., 2017). Tissues were analysed separately as they were TMT-labeled and run separately on the mass spectrometer. We considered protein contigs significantly different if they had a moderated p-value (adjusted for multiple testing) of p < 0.05 and a log fold change greater than 1. Volcano plots illustrating these cut-offs were generated with the EnhancedVolcano package (version 1.4.0, (Blighe et al., 2019)). We used a principal component analysis (PCA) to characterize variance across all proteins using the prcomp function in the R base package. We tested for group differences in principal components using the aov function in the R base package and generated PCA plots using the s.class function in the ade4 package (version 1.7–15). Heatmaps for each tissue were generated with the heatmap.2 function of the gplots package (version 3.0.3). Gene ontology enrichment analyses were performed with the R package topGO (version 2.34.0; (Rahnenfuhrer, 2019)) with significance set at p < 0.05. Boxplots were generated with the ggplot function in the ggplot2 package in R (version 3.3.0, (Wickham, 2009)).

## Results and Discussion

### Poison frogs rapidly sequester toxins

We fed decahydroquinoline (DHQ) or vehicle control to non-toxic Diablito (*Oophaga sylvatica*) poison frogs for 1, 4 or 18 days (Figure 1A) and quantified DHQ abundance across multiple tissues (Figure 1B). Although DHQ was not detectable one day after exposure, DHQ levels increased after 4 and 18 days in all tissues (Figure 1C). The abundance of DHQ varied across frog groups sampled at different times after the onset of toxin feeding (*X*^2^(1)=16.387, p=5.163×10^−5^). DHQ abundance was also different across tissues (*X*^2^(2)=10.331, p=0.001), with the highest amounts in the skin, where alkaloids are stored in granular glands. Uptake within 4 days is faster than previous studies, which documented skin alkaloid uptake after 3 months in bufonids (Hantak et al., 2013) and in 2-6 weeks in other dendrobatid species (Daly et al., 2003). We did not detect DHQ after 1 day, suggesting either DHQ did not reach the intestines by that time or abundance was below the limit of detection. Rapid uptake in dietary alkaloid toxins has important ecological implications, as acute shifts in arthropod diet could quickly contribute to the poison frog chemical repertoire and alter palatability to predators.

### Decahydroquinoline exposure induces proteome shifts across tissues

As poison frogs rapidly uptake toxins, we then asked how frog physiology changes with DHQ uptake by examining protein abundance differences between control frogs and chemically defended frogs using quantitative proteomics (Figure 2, Supplementary Excel File, Figures S3–S5). We used principal component analysis to visualize overall protein variation between groups in each tissue (Figures S3–S5). In the intestines and skin, principal component (PC) 3 separated toxic and control frogs (intestines: F(1)=31.21, p=0.0008; skin: F(1)=23.36, p=0.002) and accounted for 13% of the variance in both tissues. In the liver, we did not observe significant separation of toxic versus control frogs in the first three PCs. Some protein contigs varied significantly with toxicity, including 276 in the intestines, 185 in the liver, and 278 in the skin.

Across all tissues, there was significant gene ontology enrichment in four biological processes: negative regulation of endopeptidase activity, complement activation, inflammatory response, and oxidation-reduction processes.

**Figure 2.**
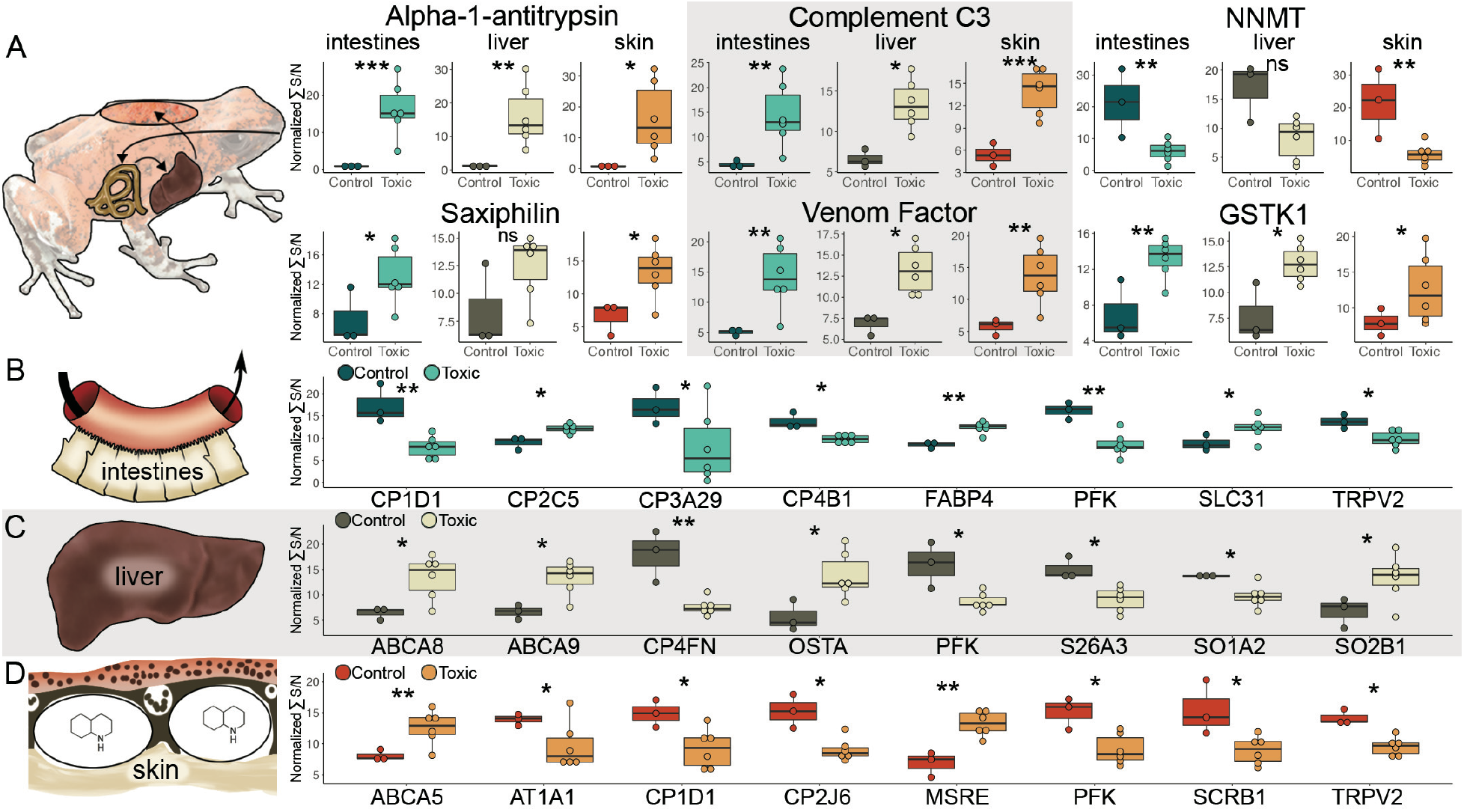
Proteomic changes with toxin bioaccumulation in the Diablito poison frog. **(A)** Some proteins changed in abundance with toxicity across all tissues, including plasma proteins alpha-1-antitrypsin, saxiphilin, complement C3, and a venom factor, as well as small molecule detoxification proteins nicotinamide N-methyltransferase (NNMT) and glutathione S-transferase kappa 1 (GSTK1). Y-axis shows the normalized sum of the signal to noise ratio. There were also many tissue-specific membrane protein abundance changes in the **(B)** intestines, **(C)** liver, and **(D)** skin that include small molecule transport and metabolism. This experiment was performed once. See Table 1 for detailed statistics (* p<0.05, ** p≤0.005, *** p≤0.0005) comparing non-toxic (N=3) and DHQ-containing (N=6) frogs. Data are represented in boxplots that show rectangles as the lower and upper quartiles (with the median as the line) and whiskers that indicate the maximum and minimum values. Abbreviations: ABCA5, ATP Binding Cassette Subfamily A Member 5; ABCA8, ATP Binding Cassette Subfamily A Member 8; ABCA9, ATP Binding Cassette Subfamily A Member 9; AT1A1, ATPase Na+/K+ Transporting Subunit Alpha 1; CP1D1, Cytochrome P450 1D1; CP2C5, Cytochrome P450 2C5; CP2J6, Cytochrome P450 2J6; CP3A29, Cytochrome P450 3A29; CP4B1, Cytochrome P450 4B1; CP4FN, Cytochrome P450 4FN; FABP4, Fatty Acid Binding Protein 4; MSRE, Macrophage Scavenger Receptor 1; OSTA, Solute Carrier Organic Anion Exchanger Family Member 26A3; PFK, Phosphofructokinase; S26A3; Solute Carrier Family 26 Member 3; SCRB1, Scavenger Receptor Class B Member 1; SLC31; Solute Carrier Family 3 Member 1; SO1A2, Solute Carrier Organic Anion Transporter Family Member 1A2; SO1B1, Solute Carrier Organic Anion Transporter Family Member 1B1; TRPV2, Transient Receptor Potential Cation Channel Subfamily V2.

Many serum proteins were differentially abundant across tissues including transferrins, globulins, complement factors, and protease inhibitors (Supplementary Excel File). Plasma protease inhibitors, which negatively regulate endopeptidase activity, were increased in abundance across all tissues, including alpha-1-antitrypsin (Figure 2A, Table 1) and alpha-2-macroglobulin. Although these proteins have been associated with rattlesnake venom resistance in squirrels (Gibbs et al., 2020), whether they bind alkaloids is currently unknown. The amphibian protein saxiphilin, which binds the lethal marine alkaloid saxitoxin (Mahar et al., 1991; Yen et al., 2019), increased in abundance in the intestines and skin of toxic frogs (Figure 2A, Table 1). We have previously shown that saxiphilin is more abundant in the plasma of non-toxic frogs and that saxiphilin may bind DHQ using thermal shift assays (Caty et al., 2019). These data support the hypothesis that saxiphilin may carry alkaloid toxins in poison frogs. Although plasma proteins may transport alkaloids through blood, binding studies are needed to determine which of these proteins bind the lipophilic alkaloids found in poison frogs.

**Table 1.** Statistics for protein contigs displayed in Figure 2. The proteins in each tissue are detailed in each row. Results are from the *limma* empirical Bayes analysis and show the log2 fold change (LogFC) the moderated test statistic (t_mod_) and the moderated p-value (p_nod_, adjusted for false discovery rate).

| Tissue | Protein | LogFC | t<sub>mod</sub> | p<sub>mod</sub> |
| --- | --- | --- | --- | --- |
| Intestines | Alpha-1-antitrypsin (A1AT) | 4.37 | 5.43 | 0.0003 |
|  | Saxiphilin (SAX) | 2.03 | 2.53 | 0.030 |
|  | Complement C3 (CO3) | 3.16 | 3.34 | 0.008 |
|  | Venom Factor (VCO3) | 2.85 | 3.96 | 0.003 |
|  | Nicotine N-methyltransferase (NNMT) | -8.69 | -3.38 | 0.007 |
|  | Glutathione S-transferase kappa 1 (GSTK1) | 1.88 | 4.03 | 0.002 |
|  | Cytochrome P450 1D1 (CP1D1) | -2.48 | -4.15 | 0.002 |
|  | Cytochrome P450 2C5 (CP2C5) | 1.06 | 3.27 | 0.008 |
|  | Cytochrome P450 3A29 (CP3A29) | -2.49 | -2.45 | 0.035 |
|  | Cytochrome P450 4B1 (CP4B1) | -1.01 | -3.07 | 0.012 |
|  | Fatty Acid Binding Protein 4 (FABP4) | 1.22 | 4.02 | 0.002 |
|  | Phosphofructokinase (PFK) | -1.9 | -4.01 | 0.002 |
|  | Solute Carrier Family 3 Member 1 (SLC31) | 1.02 | 2.72 | 0.021 |
|  | Transient Receptor Potential Cation Channel Subfamily V2 (TRPV2) | -1.02 | -2.58 | 0.027 |
| Liver | Alpha-1-antitrypsin (A1AT) | 3.49 | 2.75 | 0.020 |
|  | Complement C3 (CO3) | 1.61 | 2.98 | 0.014 |
|  | Venom Factor (VCO3) | 1.51 | 3.11 | 0.011 |
|  | Glutathione S-transferase kappa 1 (GSTK1) | 1.36 | 3.06 | 0.012 |
|  | ATP Binding Cassette Subfamily A Member 8 (ABCA8) | 1.85 | 2.48 | 0.032 |
|  | ATP Binding Cassette Subfamily A Member 9 (ABCA9) | 1.72 | 2.77 | 0.019 |
|  | Cytochrome P450 4F22 (CP4FN) | -2.55 | -4.82 | 0.0007 |
|  | Organic Solute Transporter Subunit Alpha (OSTA) | 2.15 | 2.86 | 0.017 |
|  | Phosphofructokinase (PFK) | -1.81 | -3.44 | 0.006 |
|  | Solute Carrier Organic Anion Exchanger Family Member 26A3 (S26A3) | -1.49 | -2.84 | 0.017 |
|  | Solute Carrier Organic Anion Transporter Family Member 1A2 (SO1A2) | -1.02 | -2.41 | 0.036 |
|  | Solute Carrier Organic Anion Transporter Family Member 2B1 (SO2B1) | 1.78 | 2.38 | 0.038 |
| Skin | Alpha-1-antitrypsin (A1AT) | 4.84 | 2.72 | 0.021 |
|  | Saxiphilin (SAX) | 2.31 | 3.14 | 0.01 |
|  | Complement C3 (CO3) | 2.72 | 5.02 | 0.0005 |
|  | Venom Factor (VCO3) | 2.58 | 3.60 | 0.005 |
|  | ATP Binding Cassette Subfamily A Member 5 (ABCA5) | 1.55 | 3.32 | 0.008 |
|  | ATPase Na<sup>+</sup>/K<sup>+</sup> Transporting Subunit Alpha 1 (AT1A1) | -1.14 | -2.33 | 0.042 |
|  | Cytochrome P450 1D1 (CP1D1) | -1.41 | -2.24 | -0.049 |
|  | Cytochrome P450 2J6 (CP2J6) | -1.54 | -3.29 | 0.008 |
|  | Macrophage Scavenger Receptor 1 (MSRE) | 1.87 | 4.75 | 0.0007 |
|  | Phosphofructokinase (PFK) | -1.46 | -2.99 | 0.013 |
|  | SCRB1, Scavenger Receptor Class B Member 1 | -1.66 | -2.88 | 0.016 |
|  | Transient Receptor Potential Cation Channel Subfamily V2 (TRPV2) | -1.21 | -3.47 | 0.006 |

A strong pattern observed in this study was that the complement system, which plays a role in adaptive and innate immunity (Sahu and Lambris, 2001), is more active in toxic frogs (Figure 2A). Complement C3 (CO3) and a related protein, A.superbus venom factor, were increased in toxic frogs in all tissues (Table 1). Complement C3 has been linked to poison frog toxicity previously through RNA sequencing studies (Caty et al., 2019; Sanchez et al., 2019), but this is the first proteomic evidence across tissues. The significant upregulation of a complement C3-like venom factor was particularly interesting. Venom complement factors were first discovered in cobras and later identified in other venomous animals, like spiders (Tambourgi and van den Berg, 2014). Cobra Venom Factor (CVF)-like proteins are co-injected with toxins from the venom glands, but are not toxic in themselves. Rather, CVF-like proteins activate complement C3 and deplete the complement system in the victim, which increases vascular permeability at the envenomation site, spreading the toxins throughout the prey faster (Vogel and Fritzinger, 2010). Although the function of CVF-like proteins in amphibians is unknown, they could have alkaloid absorption enhancing capabilities, as has been suggested for other amphibian skin peptides (Raaymakers et al., 2017), but further testing is necessary to establish this as a complement-dependent phenomenon.

Across all tissues, a number of membrane proteins involved in drug metabolism were differentially abundant between toxic and control frogs. One of the strongest protein responses to toxicity was a decrease of nicotinamide N-methyltransferase abundance in the intestines and skin of toxic frogs (Figure 2A). This protein detoxifies xenobiotics by methylating the nitrogen in pyridine rings (Alston and Abeles, 1988). Similarly, many cytochrome P450s, well known for their small molecule metabolism (Danielson, 2002), decreased in abundance with toxicity (Figures 2B-D, S2). These patterns were surprising, as we initially hypothesized that small molecule metabolism capacity would increase with toxin accumulation, and indeed there are some cytochrome P450s that increase with DHQ accumulation in the skin (Figure S5). However, this decrease in abundance of proteins that metabolize small molecules may instead reflect physiological reduction in xenobiotic metabolism so that toxins can arrive intact to the skin. On the other hand, glutathione S-transferase kappa 1, which is involved in cellular detoxification by conjugating glutathione to hydrophobic substances for export, increased in toxic frogs in the intestines and skin. Acquiring toxins likely involves a delicate balance between storing the compound with intact potency and avoiding autotoxicity. Determining which of these enzymes modifies poison frog alkaloids would be a useful step towards understanding this physiological balance.

Abundance of many membrane transporter proteins changed with DHQ accumulation, including ion channels, ABC (ATP-binding cassette) transporters (Schinkel and Jonker, 2003) and solute carrier proteins (Lin et al., 2015). The transient receptor potential cation channel subfamily V member 2 (TRPV2) is involved in the detection of noxious chemicals (Perálvarez-Marín et al., 2013) and decreased in abundance in the intestines and skin of toxic frogs. We also note that Na+/K+-ATPase is downregulated with toxin accumulation in the skin. Mutations in this protein are associated with toxin-resistance mechanisms in many animals (Ujvari et al., 2015) and changing the abundance of this protein may be an additional coping mechanism. Additionally, ABC and solute carrier proteins bind to xenobiotics in mammals, including in the context of cancer resistance to alkaloid-based anticancer therapies (Chen et al., 2016; Liu, 2009). These proteins may be important in uptake and transfer of alkaloids across tissues, as we found changes in abundance of several proteins in this family. For example, the increase in abundance of the neutral and basic amino acid transport protein (SLC31) in the intestines contributed to the separation of toxic and control frogs in the PCA. More in depth *in vitro* studies with these membrane transporter proteins would be required to fully understand these trends and the extent to which these proteins can interact with poison frog alkaloids.

Focusing on proteins that contribute to PCA separation of control and toxic frogs, we found an elevation of lipid-associated processes and a decrease in some glycolysis-related proteins, suggesting acquiring toxins shifts cellular metabolism more towards aerobic than anaerobic processes. Specifically, ATP-dependent 6-phosphofructokinase (PFK), the enzyme for the first committing step in glycolysis, is decreased in all tissues in toxic frogs, while many proteins involved in lipid transport and metabolism are increased. Chemically defended dendrobatid poison frogs have greater aerobic capacity (Santos and Cannatella, 2011) compared to non-toxic species and this study provides a potential physiological explanation for this observation. Lipid-associated proteins also point to an alternative mechanism of alkaloid transport, where lipophilic compounds can be transported via chylomicrons in the lymphatic system (Trevaskis et al., 2008). For example, the fatty acid binding protein, which transports lipophilic substances, increases in the intestines of toxic frogs (Figure 2B, Table 1). Scavenger receptor proteins involved in lipoprotein endocytosis also change in abundance in the skin of toxic frogs and provide a potential sequestration mechanism. For lipid-associated transport to be feasible, proteins that break down lipids are also needed, and indeed, lipases are also increased in the skin of toxic frogs. It is currently an open question as to whether the main route of alkaloid uptake in poison frogs is carrier-mediated through blood circulation or lipid-mediated transport through the lymph system.

In this study, we show that the Diablito frog can bioaccumulate dietary alkaloids within four days of exposure, and this rapid uptake shifts abundance of proteins that interact with xenobiotics, including plasma carrier proteins, membrane transporters, cytochrome P450s, and lipoproteins. Our study also suggests that the complement system and venom factors may be involved in alkaloid delivery, although experiments involving frog predators are needed to test this idea. Our study lays a foundation for future work involving rigorous small molecule binding, transport, and metabolism experiments, which are needed to pinpoint specific physiological changes that allow poison frogs to acquire their chemical defenses.

## Acknowledgements

We thank Aurora Alvarez-Bullya, Stephanie Caty, and Nora Moskowitz for comments on early versions of this manuscript and Eva Fischer and Alexandre Roland for teaching assistance in the LS50 laboratory course. We acknowledge that our research at Stanford University occurs on the ancestral and unceded land of the Muwekma Ohlone Tribe.

## Data availability

Mass spectrometry data are available at PRIDE database (PRoteomics IDEntifications Database, Project accession PXD021216). All processed data is in the Supplementary Excel File.

## Author Contributions

LAO and AWM designed the research; Harvard College students (LS50:Integrated Science: Marianne T. Aguilar, Sophia M. Caldera, Jacqueline Chea, Miruna G. Cristus, Jett P. Crowdis, Bluyé DeMessie, Caroline R. DesJardins-Park, Audrey H. Effenberger, Felipe Flores, Michael Giles, Emma Y. He, Nike S. Izmaylov, ChangWon C. Lee, Nicholas A. Pagel, Krystal K. Phu, Leah U. Rosen, Danielle A. Seda, Yong Shen, Santiago Vargas, and Hadley S. Weiss) funded by the Howard Hughes Medical Institute Professor’s Award 520008146 (AWM) conducted the toxin feeding experiment and preliminary data analyses under the guidance of LAO; JDO and JAP conducted the proteomics experiments; SAT quantified DHQ; SPG and AWM contributed new reagents/tools; LAO analyzed the data and wrote the paper with contributions from all authors.

## Supplementary Figures

**Figure S1.**
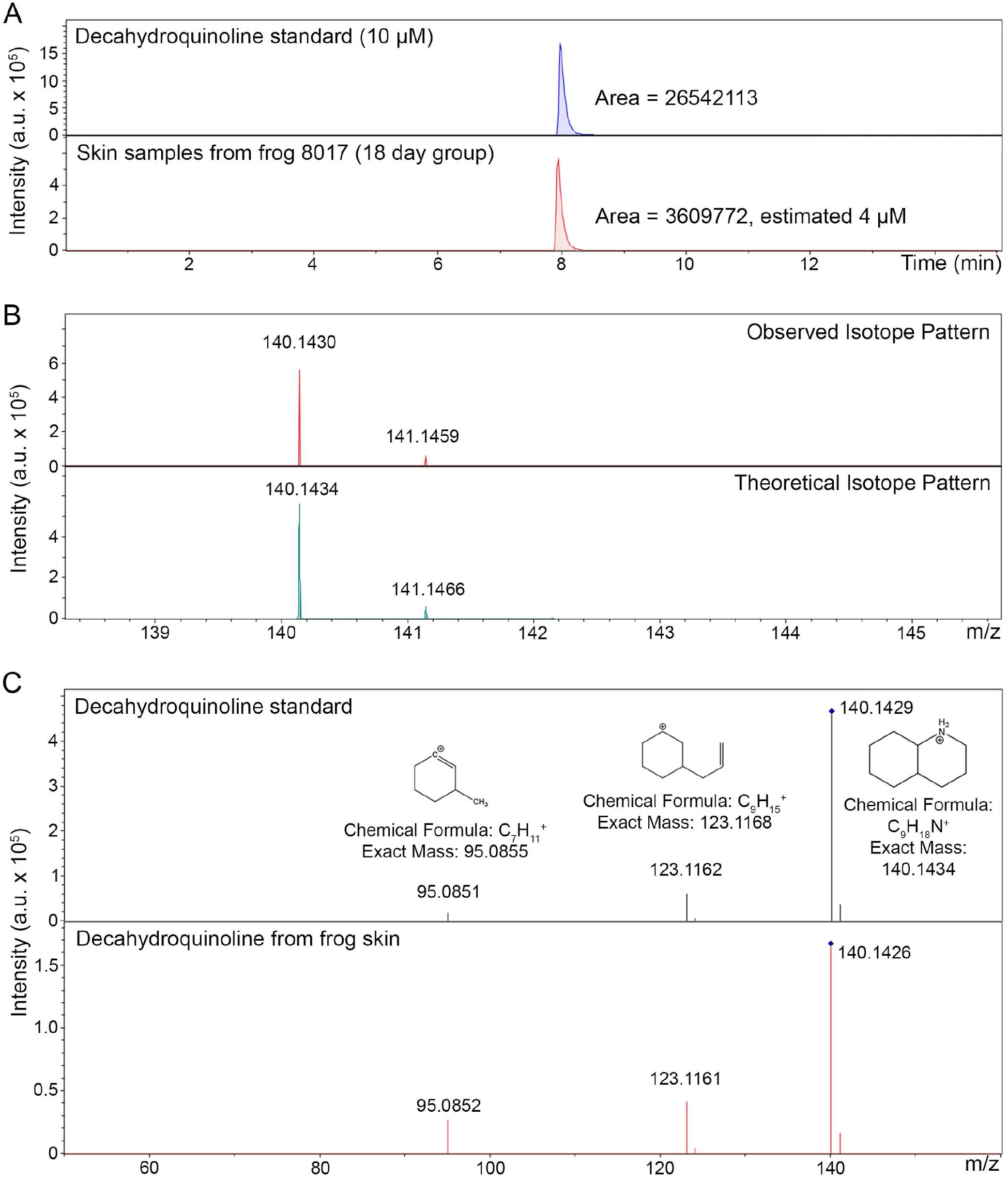
Detection of decahydroquinoline (DHQ) using liquid chromatography / mass spectrometry. **(A)** DHQ was dissolved at 10μM in methanol and used as a standard (top panel) to quantify tissue samples (bottom panel). **(B)** The observed isotope pattern of the DHQ standard (top panel) matched the theoretical expectation (bottom panel). **(C)** Tandem mass spectrometry was used to confirm the structure of DHQ from frog skin with the standard.

**Figure S2.**
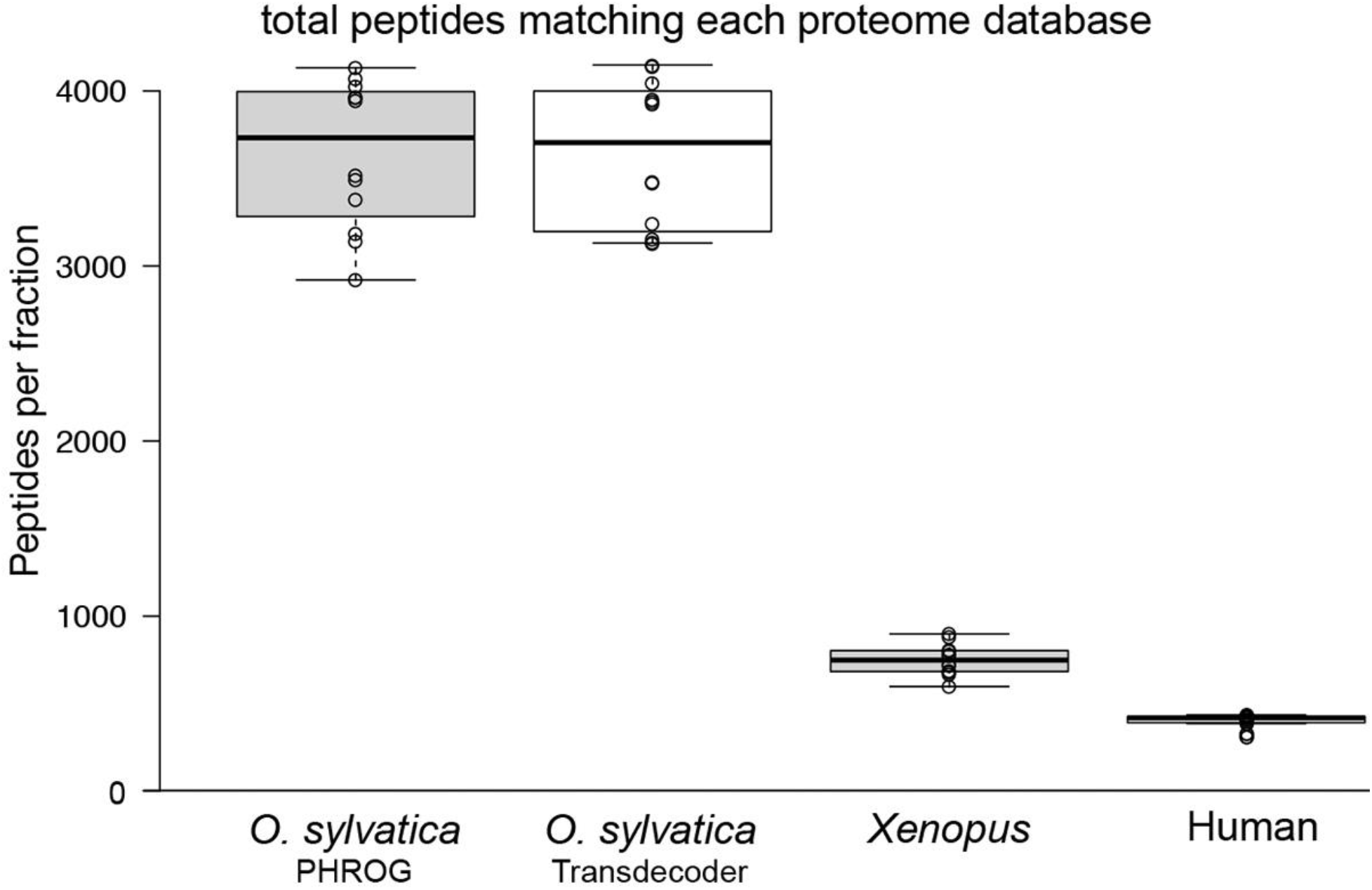
Comparison of proteome databases in peptide matching. The number of matching peptides from all tissues in the DHQ feeding dataset was compared across different proteome references. Two *O. sylvatica* references were equivalent, including one reference generated using the PHROG workflow (proteomic reference with heterogeneous RNA omitting the genome; (Wühr et al., 2014)) and the other generated using Transdecoder (http://**transdecoder**.github.io). We also compared the reference proteomes for *Xenopus* and human, which had fewer matches than the species-specific databases generated from transcriptome data. The *O. sylvatica* PHROG reference proteome was used in the final analysis.

**Figure S3.**
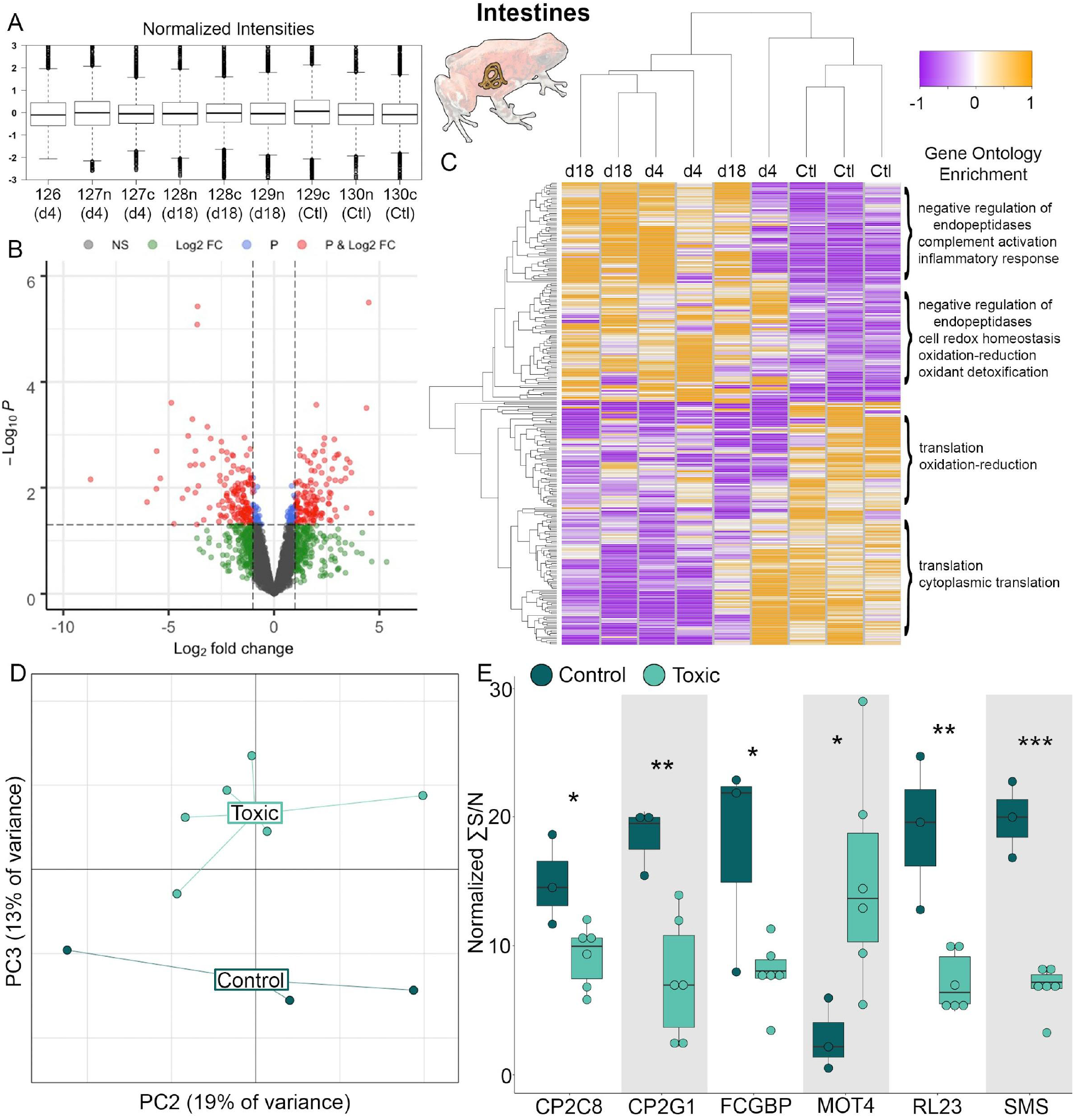
Quantification of protein changes in the intestines with decahydroquinoline bioaccumulation. **(A)** Protein samples were tandem mass tag (TMT) labeled prior to quantification and channels were z-score normalized prior to analysis. **(B)** Volcano plots show protein contigs that are not significant (NS, black), log fold change greater than one (Log2 FC, green), a p-value < 0.05 (P, blue) or both (red). **(C)** Heatmap of protein contig abundance (rows) between individuals frogs (columns); gene ontology enrichments in different clusters are on the right. **(D)** Visualization of a principal component analysis, where PC3 significantly separated control and toxic frogs (p=0.0008). **(E)** Examples of proteins that are significantly different between non-toxic (N=3) and DHQ-containing (N=6) frogs (* p<0.05, ** p≤0.005, *** p≤0.0005). Data are represented in boxplots that show rectangles as the lower and upper quartiles (with the median as the line) and whiskers that indicate the maximum and minimum values. Abbreviations: CP2C8, Cytochrome P450 Family 2 Subfamily C Member 8; CP2G1, Cytochrome P450 Family 2 Subfamily G Member 1; FCGBP, IgG Fc Binding Protein; MOT4, Monocarboxylate Transporter 4; RL23, Ribosomal Protein L23; SMS, Somatostatin.

**Figure S4.**
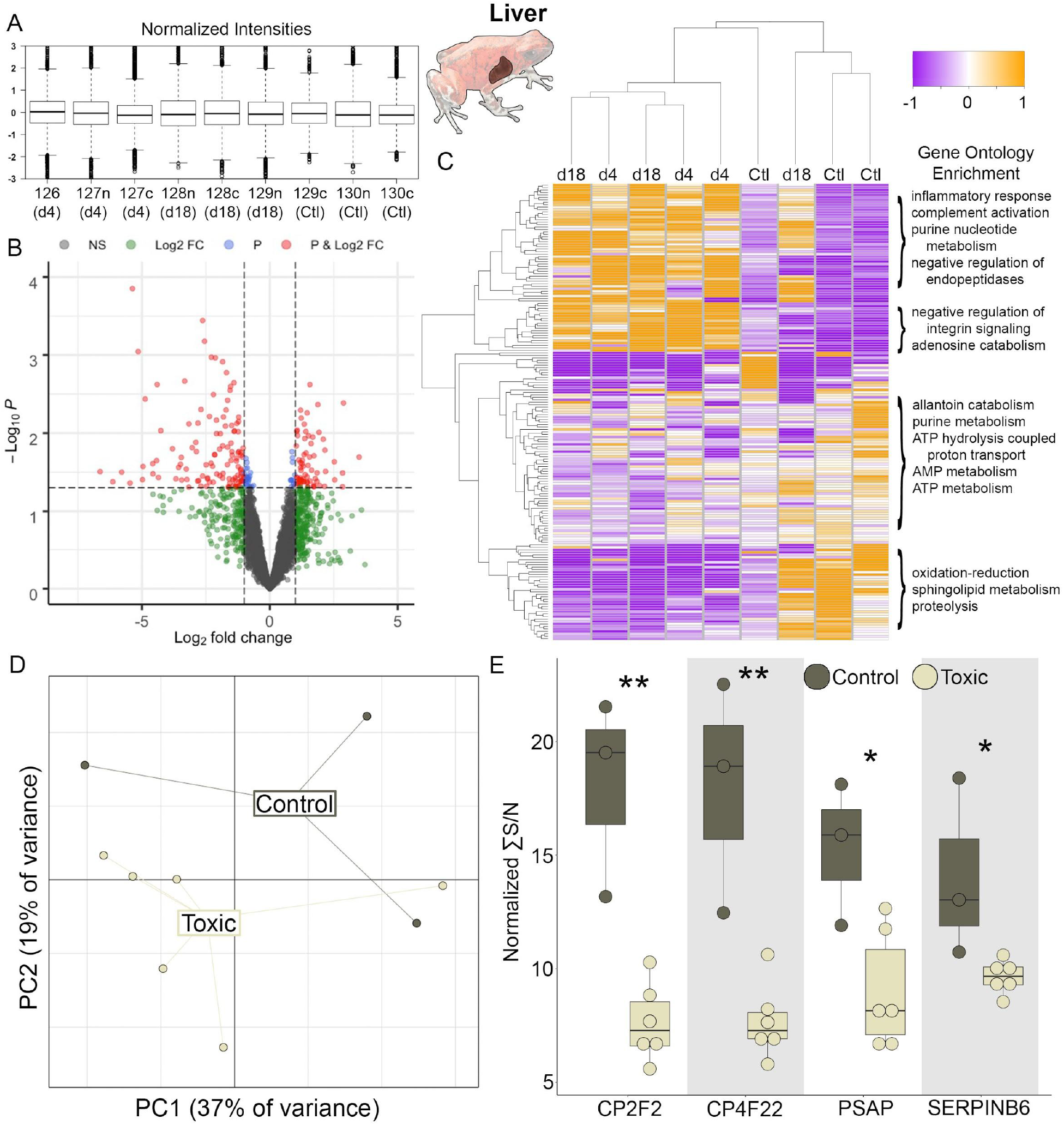
Quantification of protein changes in the liver with decahydroquinoline bioaccumulation. **(A)** Protein samples were tandem mass tag (TMT) labeled prior to quantification and channels were z-score normalized prior to analysis. **(B)** A volcano plot shows protein contigs that are not significant (NS, black), log fold change greater than one (Log2 FC, green), a p-value < 0.05 (P, blue) or both (red). **(C)** A heatmap of protein contig abundance (rows) between individuals frogs (columns); gene ontology enrichments in different clusters are on the right. **(D)** Visualization of a principal component analysis; groups were not significantly separated in the first three principal components. **(E)** Examples of proteins that are significantly different between non-toxic (N=3) and DHQ-containing (N=6) frogs (* p<0.05, ** p≤0.005, *** p≤0.0005). Data are represented in boxplots that show rectangles as the lower and upper quartiles (with the median as the line) and whiskers that indicate the maximum and minimum values. Abbreviations: CP2F2, Cytochrome P450 Family 2 Subfamily F Member 2; CP4F22, Cytochrome P450 Family 2 Subfamily F Member 22; PSAP, Prosaposin; SERPINB6, Serpin P6.

**Figure S5.**
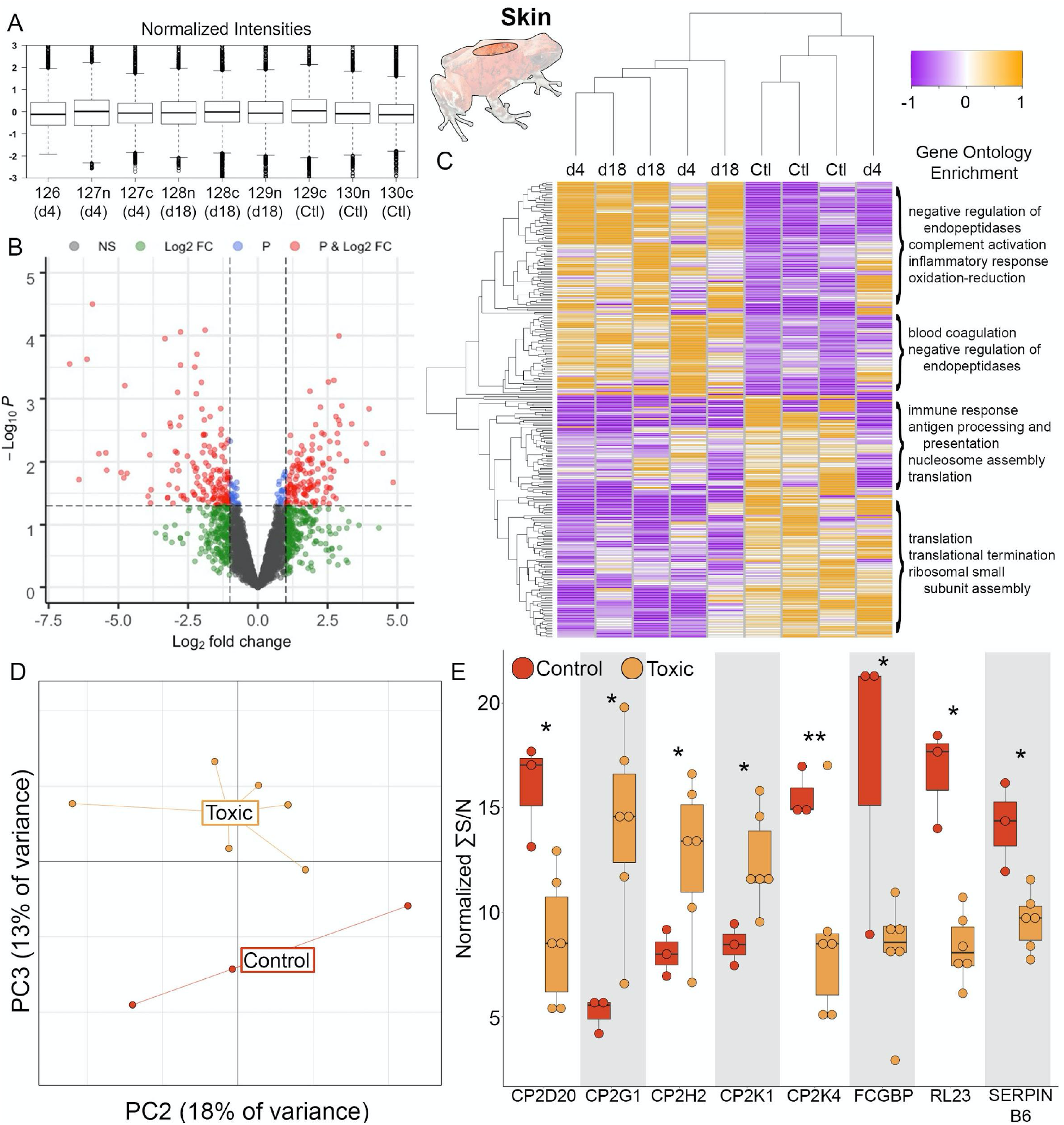
Quantification of protein changes in the skin with decahydroquinoline bioaccumulation. **(A)** Protein samples were tandem mass tag (TMT) labeled prior to quantification using mass spectrometry. Channels were z-score normalized prior to analysis. **(B)** A volcano plot shows protein contigs that are not significant (NS, black), log fold change greater than one (Log2 FC, green), a p-value < 0.05 (P, blue) or both (red). **(C)** A heatmap of protein contig abundance (rows) between individuals frogs (columns); gene ontology enrichments in different clusters are on the right. **(D)** Visualization of a principal component analysis, where PC3 significantly separated control and toxic frogs (p=0.002). **(E)** Examples of proteins that are significantly different between non-toxic (N=3) and DHQ-containing (N=6) frogs (* p<0.05, ** p≤0.005, *** p≤0.0005). Data are represented in boxplots that show rectangles as the lower and upper quartiles (with the median as the line) and whiskers that indicate the maximum and minimum values. Abbreviations: CP2D20, Cytochrome P450 Family 2 Subfamily D Member 20; CP2G1, Cytochrome P450 Family 2 Subfamily G Member 1; CP2H2, Cytochrome P450 Family 2 Subfamily H Member 2; CP2K1, Cytochrome P450 Family 2 Subfamily K Member 1; CP2K4, Cytochrome P450 Family 2 Subfamily K Member 4; FCGBP, IgG Fc Binding Protein; RL23, Ribosomal Protein L23.

